# SILAC-based quantitative proteomic analysis of *Drosophila* gastrula stage embryos mutant for fibroblast growth factor signaling

**DOI:** 10.1101/707232

**Authors:** Hamze Beati, Alistair Langlands, Sara ’ten Have, H.-Arno J. Müller

**Affiliations:** Developmental Genetics Unit, Institute of Biology, University of Kassel, Germany; Division of Cell and Developmental Biology, School of Life Sciences, University of Dundee, United Kingdom; Division of Gene Regulation and Expression, School of Life Sciences, University of Dundee, United Kingdom

**Keywords:** Drosophila, SILAC, Fibroblast Growth Factor, Cell Signaling, proteomics

## Abstract

The application of quantitative proteomics in model organisms has been successful in determining changes in the proteome under distinct physiological conditions. Quantitative mass spectrometry-based proteomic analyses in combination with genetics provide powerful tools in developmental cell signaling research. *Drosophila melanogaster* is one of the most widely used genetic models for studying development and disease. Here we combined quantitative proteomics with genetic selection to determine global changes in the proteome upon depletion of the Heartless (Htl) Fibroblast-Growth Factor (FGF) receptor signaling in *Drosophila* embryos at early gastrulation stages. We present a robust, single generation SILAC (stable isotope labeling with amino acids in cell culture) protocol for labeling proteins in early embryos and for selection of homozygously mutant embryos at pre-gastrula stages using an independent genetic marker. Our analyses detected quantitative changes in the global proteome of *htl* mutant embryos during gastrulation. We identified distinct classes of down-regulated and up-regulated proteins and network analyses indicates functionally related groups of proteins in each class. These data suggest that FGF signaling in the early embryo affects global changes in the abundance of metabolic, nucleoplasmic, cytoskeletal and endomembrane transport proteins.

## Introduction

Quantitative mass spectrometry-based proteomics have been implemented in studying levels of protein expression and protein modifications in various cell types, tissues and organisms for comparing diverse states, such as age, gender, drug treatments or various disease conditions. In 2002, the SILAC (<u>s</u>table isotope labeling with <u>a</u>mino <u>a</u>cids in <u>c</u>ell culture) method was introduced as a tool for studying functional proteomics on a quantitative global scale^1^. SILAC-based mass spectrometry allows to directly compare populations of cells on the basis of differential signature labeling with stable isotopes. For example, one cell line grown on media containing the naturally predominant occurring amino acid is compared with cells grown on media containing a stable isotope-labeled form of an amino acid ^2^. SILAC labeling has also proven successful for global quantitative proteomic comparisons of entire organisms including mammalian model systems^3^. Since then various multicellular model organisms were successfully labeled with stable non-radioactive isotopes (SILAC or ^15^N labeling) including the plant *Arabidopsis thaliana*, the nematode *Caenorhabditis elegans*, the fly *Drosophila melanogaster* and the mammal *Mus musculus*^4-13^.

*Drosophila melanogaster* is one of the most widely studied genetically tractable model organism and has been employed for more than a century to improve our understanding in many areas in biology including genetics, developmental cell biology and signal transduction. Protocols for SILAC or ^15^N labeling of *Drosophila* are based on feeding flies with yeast that has been labeled with stable isotopes^4, 9, 11^. Stable isotope labeling in flies has been used to determine sex-specific differences in the proteome of somatic cells^9^, differences in the proteome during ageing^14^, and differences in proteomes between adults, larvae and pupae^15, 16^. Recently, a quantitative proteomic study investigated the developmental profile of the *Drosophila* proteome throughout the life cycle using label-free approaches^15^. Proteome studies in embryos obtained from SILAC flies or using label-free methods also addressed proteome dynamics in early development like changes during the oocyte-to-egg transition (oocyte maturation), and the alterations that occur during the transition of the maternal to the zygotic transcriptional program (maternal/zygotic transition)^8, 17, 18^. Despite the advances in quantitative proteomics and the plethora of mutations in developmental control genes, changes in the proteome of fly embryos homozygously mutant for a recessive developmental allele have not yet been performed due to technical difficulties in obtaining sufficient material.

In this study we combined the power of *Drosophila* genetics with SILAC labeling to examine changes that occur in the proteome when a major developmental signaling pathway is lacking in the early embryo. The *Drosophila* gastrula stage embryo contains a relatively low cellular complexity, but the cells participate in major morphogenetic movements^19^. Most dramatically, the prospective mesoderm germ layer moves into the interior of the embryo and undergoes an epithelial-to-mesenchymal transition (EMT) and collective cell migration. These morphogenetic events are controlled by the activation of a fibroblast growth factor (FGF) receptor encoded by the *heartless* (*htl*) gene ^20-26^. We emloss-of-function mutations in *htl* to investigate the effects of FGFR signaling in the context of an entire embryo. The response of mesoderm cells upon Htl receptor activation is rapid as the cells form cellular protrusions and move towards the underlying ectoderm within the range of minutes ^27^. Therefore, it has been suggested that the Htl signal initially affects posttranslational modifications and the synthesis and turnover of proteins involved in cell movements rather than transcriptional responses ^27, 28^.

One major problem that hampered the comparative analyses of proteomes from wild-type and mutant embryos was the selection of the homozygously mutant embryos, which make only 25% of the progeny from heterozygous parental animals. A phenotypic selection marker that serves this purpose should be detectable early in development, ideally before the gene under investigation becomes active and its mutant phenotype becomes visible. Furthermore, the marker should not affect the viability of the embryo or the organism. We established a system using the *halo* mutation which allows the selection of homozygous mutant embryos in early developmental stages^29^. The *halo* mutation causes a readily visible defect in the transport of lipid droplets in early embryos, but does not affect viability and fertility ^29,30^. Here we present a protocol for efficient labeling of *Drosophila* embryos with stable non-radioactive isotopes in a single generation combined with genotyping and staging early embryos to discriminate between homozygous *htl* mutant embryos and *htl* heterozygous embryos (Fig. 1A,B). Quantitative global proteomic analysis of *htl* mutant embryos resulted in the discovery of protein networks that were down- or up-regulated when compared to control embryos.

**Figure 1:**
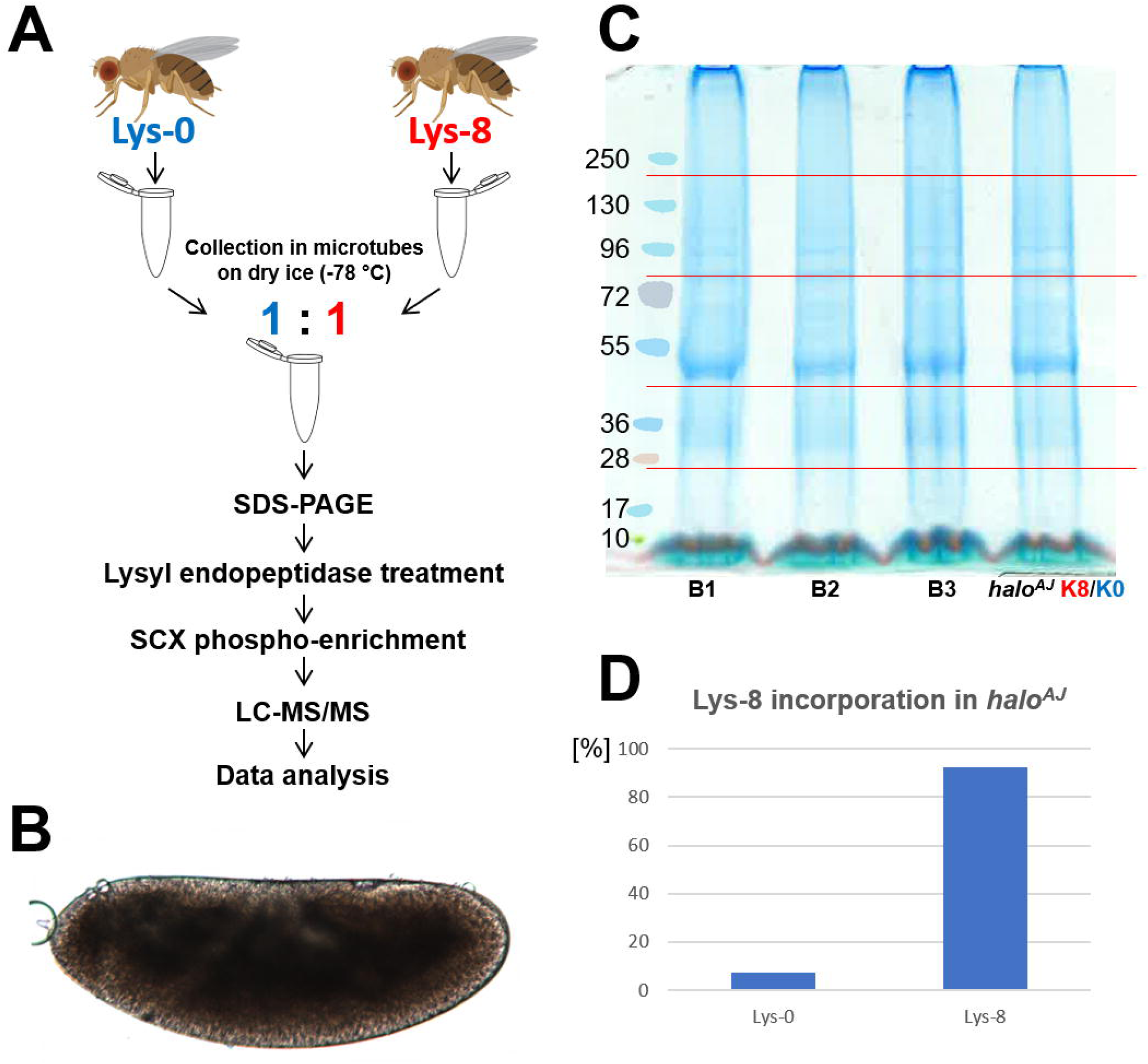
SILAC workflow using *Drosophila* embryos. **(A)** *Drosophila* flies were grown on media containing Lys-0 or Lys-8 labeled yeast, respectively and embryos were collected in microtubes on dry ice. After embryo lysis equal amounts of protein were mixed, separated via SDS-PAGE and cleaved into peptides with a Lys-C endopeptidase. Following SCX chromatography, samples were run on a *LC-MS/MS Velo Orbitrap* instrument and data analysis was carried out with MaxQuant and Perseus. **(B)** *Drosophila* embryos were collected during early stage 8. Anterior is left, dorsal up. **(C)** 3 biological replicates (B1-B3) and a Lys-8 *halo*^*AJ-/-*^ / Lys-0 *halo*^*AJ-/-*^ comparison are shown after SDS-PAGE and Coomassie staining. Red lines indicate separation of the sample lines for further preparation. **(D)** Rearing *halo*^*AJ*^ embryos on Lys-8 containing yeast leads to heavy labeled flies that produce embryos with a Lys-8 incorporation of > 92%.

## Methods

### *Drosophila* strains

Fly stocks were kept under standard conditions. The stock containing the loss of function *halo*^*AJ*^ allele and a stock harboring a transgene with the *halo* genomic locus (*p[halo]*) were gifts of M. Welte (Univ. of Rochester, U.S.A.)^31, 32^. We used a chromosome harboring transposase Δ2,3 under the control of a hsp70 promoter to mobilize the *p[halo]* transposon insertion in the genome and isolated insertions on the autosomal balancer chromosomes TM3, TM6 and CyO. The loss-of-function *htl*^*AB42*^ allele was maintained with a TM6B, *Hu, Tb, e, p[halo]* balancer chromosome and crossed into a homozygous *halo*^*AJ*^ background.

### SILAC labeling of *Saccharomyces cerevisiae*

*Saccharomyces cerevisiae BY4742* colonies were allowed to grow for 2 days at 30°C. A single colony was inoculated into 5 ml DOA (Dropout) – no Lysine media (synthetic complete) supplemented with 5 μl of heavy Lysine (Lys-8; stock: 30 mg/ml) (L-Lysine: 2HCl, U-^13^C_6_, 99%; U-^15^N_2_, 99%, Cambridge Isotope Laboratories, Inc.). The culture was incubated at 30°C for 24 hours. 5 μl of cell suspension was used to inoculate 5 ml of fresh media containing Lys-8 and was incubated for another 24 hours at 30°C in a shaking incubator. 1 ml of culture was used to inoculate 1 l of DOA media containing Lys-8. Incubation took place for another 24 hours at 30°C in a shaking incubator. 1 ml of the culture was saved for a label check and the remaining culture was pelleted by centrifugation. The yeast pellet was resuspended once in dH_2_O and centrifuged; the supernatant was discarded and the Lys-8 – labeled yeast was stored at −80°C. Previous protocols used the Lysine auxotrophic *S. cerevisiae* stock *SUB62* ^9^. We found that the *SUB62* was insufficiently labeled in a Lys-8 containing medium with an average global SILAC ratio of 1.4 and below suggesting a maximum of 58.4 % Lys-8 label (Suppl. Fig. S1A,B). *SUB62* strain harbors the point mutation *lys2-801*, which was described as an amber mutation in the *lys2* gene and therefore is in principle revertible in particular when grown in large cultures. We found that in 1 l cultures the *SUB62* strain undergoes reversion to a lys prototrophic strain and that this effect appears to be enhanced in the presence of Lys-8 as a source (Suppl. Fig. S1). We therefore utilized the strain *BY4742* for our experiments, which contains the lys2Δ0 mutation, which is a complete deletion of the *lys2* gene and does not undergo reversions^33^. Labeling effiency with *BY4742* was nearly complete exhibiting a global SILAC ratio of around 15 (Suppl. Fig. S1C,D).

### SILAC labeling *of Drosophila melanogaster*

150 embryos were transferred onto a fresh apple juice agar plate supplemented with 300 µl of *BY4742* Lys-8 yeast and enclosed with a fly cage. Larvae were allowed to hatch and to feed on Lys-8 labeled yeast at 25°C. Once larvae started to penetrate the apple juice agarose dH_2_O was supplemented according to humidity and evaporation to keep apple juice agarose/yeast mixture soft and moist. Pupae were gently transferred onto fresh apple juice agar plate supplemented with a drop of *BY4742* Lys-8 yeast to serve as food for hatching SILAC flies. Control flies of the respective genotype were raised according to the same protocol except for using yeast grown on standard Lys-0 containing media.

### Protein extraction from yeast

1 ml of Lys-8 *BY4742* culture was centrifuged, supernatant discarded and yeast pellet resuspended in 150 µl of 2M NaOH, 1 M β-Mercaptoethanol. 5 volumes of protein extract were supplemented with 1 volume of 20 % Trichloracetic acid (TCA) and mixed by inversion. The mixture was incubated on ice for 10 minutes and subsequently centrifuged at 14.000 rpm for 4 minutes at 4°C. The supernatant was removed, pellet washed with cold acetone and air dried. The pellet was resuspended in 8 M urea, 0.4 M ammonium bicarbonate and used for label check via mass spectrometry.

### Embryo collection and protein lysis

The embryos were incubated on a agarose coated petri dish, covered with halo carbon oil (27S) and were staged under a dissection microscope in transmitted light. Embryos at the desired stages were dechorionated in 0.5% sodium hypochlorite solution and collected in microtubes (Protein LoBind tubes, Eppendorf), which were cooled on dry ice. For lysis, RIPA buffer (50 mM Tris-HCl pH 8.0, 150 mM NaCl, 1 % NP40, 0.5 % Sodiumdesoxycholate, 0.1 % SDS) was supplemented with proteinase inhibitors (EDTA-free Protease Inhibitor Cocktail (complete), Roche), phosphatase inhibitors (PhosSTOP, Roche); 100 μl of buffer was used for embryo lysis for each individual biological replicate. An equal number of embryos were homogenized with a bio-vortexer and incubated on ice for 20 minutes, followed by a centrifugation step for 10 minutes at 14.000 rpm at 4°C. The supernatant was transferred into a fresh microtube and protein concentration was determined (Peptide and Protein Quantification Kit, LavaPep).

### In-gel digestion

For each biological replicate, 40 μg of protein were run on SDS-PAGE (4-12% Bis Tris Nupage gels). Each lane was cut into 5 gel pieces for an in-gel digestion (Fig. 1C), and every individual piece was cut into smaller fragments that were rinsed with 100 μl 100 mM NH_4_HCO_3_ : 100% acetonitrile (ACN) for 10 minutes at room temperature in a shaking incubator. The solution was removed from the gel pieces and the washing step was repeated. 50 μl of 100% ACN were added until gel pieces formed an aggregate and turned white. 50 μl of 100 mM NH_4_CO_3_ were added and incubated at 37°C for 30 minutes, while shaking. The solution was removed, and gel pieces were dried in a vacuum centrifuge. 50 μl of 10 mM DTT were added and incubated at 55°C for 45 minutes. After removing DTT, 50 μl of 50 mM iodoacetamide solution was added and the gel pieces incubated for 30 minutes in the dark at room temperature. After removing the iodoacetamide solution, gel pieces were washed again twice with 100 μl 100 mM NH_4_HCO_3_ : 100% ACN for 10 minutes at room temperature in an shaking incubator. After removing the washing solution, gel pieces were dried in a vacuum centrifuge. In gel digestion with the Lysyl endopeptidase (Lys-C) was performed overnight according to the manufacturers protocol (Lysyl Endopeptidase, Mass Spectrometry Grade, Fujifilm Wako Pure Chemical Corporation). After digestion, 20 μl of 0.1% TFA and 20 μl 100% ACN were added and the mixture sonicated in an ice water bath for 15 minutes. The supernatant was transferred into a new microtube and 100 μl 30% ACN : 0.1% TFA were added to the gel pieces and sonicated in an ice water bath as before. The supernatant was transferred to previously collected supernatant, and 100 μl 50% ACN : 0.1% TFA were added to the gel pieces and handled as before. Pooled supernatant was finally vacuum centrifuged at 60°C to reduce the volume to approximately 100 μl. The peptides were further cleaned up with C18 columns using HPLC according to standard protocols (GRE support group University of Dundee, Scotland).

### SCX chromatography

The sample has been reconstituted in 500 μl SCX loading/wash buffer (10 mM KH_2_PO_4_, 25% ACN, pH 3). SCX columns (Thermo Scientific Hypersep SCX; benzosulfonic acid, 25 mg/ml) were washed twice with 1 ml MilliQ water and were primed twice with 1 ml SCX priming/elution buffer (10 mM KH_2_PO_4_, 25% ACN, 350 mM KCl, pH 3). Sample was loaded onto the column and pushed through slowly. Loaded columns were washed twice with 500 μl SCX wash buffer. Sample elution was achieved with 500 μl of SCX elution buffer and followed by clean up with C18 columns.

### LC-MS/MS analysis

The peptide samples were run on a Thermo Fischer Orbitrap Velos Pro. They were separated on an Easy- Spray reversed chromatography C18 Column (ES803A, 75 μm, 500 mm) on a 150 min. gradient (buffer A: 0.1% formic acid, buffer B: 80% acetonitrile/0.1% formic Acid). The resolution of the first MS run was 60,000 (scanning from 335-1800 m/z), with top 16 ions selected for MS run 2.

### Data analysis

Raw MS data were analyzed with MaxQuant ^34^ version 1.5.2.8 and searched against the Uniprot *Drosophila* database. A range of variable modifications that were detected included acetylation (amino-terminal), oxidation (M), deamidation (NQ), Gln->pyro-Glu, Phospho (STY), and the fixed modification carbamidomethylation (C). Protein and peptide False Discovery Rate (FDR) cut offs were both set to 0.01, and a minimum peptide length was set to 7 amino acids. Statistical analysis was carried out with Perseus^35^ and Microsoft Excel. Protein network analyses of down- and up-regulated proteins were conducted with STRING version 10.5 (https://string-db.org/) ^36^.

## Results

### Generation of large quantities of SILAC flies

The aim of this study was to determine changes in the proteome of tightly staged gastrula embryos that are depleted of signaling through the FGF receptor Htl. The collection of tightly staged homozygous *htl* mutant embryos requires large quantities of heterozygous flies, because only a quarter of the embryos of this fly stock will be homozygous for the mutation and the time window for collection only lasts about 15 min. The SILAC fly was described previously using protocols that reared flies on minimal media, e.g. Lys-8 labeled yeast on cotton wool with sucrose or low-melt agarose containing glucose for efficient labeling ^9, 11^. In our hands these procedures did not produce enough healthy flies; larvae developed initially normal and formed pupae, but many flies died before eclosion. In order to obtain a larger amount of SILAC flies required for collecting sufficient quantities of staged embryos, we set out to improve the labeling procedure (see methods). In particular, substitution of the agarose with apple juice agar did not compromise the labeling efficiency with Lys-8 and produced large quantities of healty SILAC flies labeled with stable isotopes that produce embryos with >92% labeling efficiency (Fig. 1D).

### Identification of pre-gastrula stage *htl* mutant embryos

Before gastrulation commences, the *Drosophila* embryo consists of a monolayered epithelium, called blastoderm epithelium, that surrounds a central yolk cell^37, 38^. The mesoderm germ layer originates from the ventral domain of the blastoderm epithelium and is internalized in a process called mesoderm invagination. After mesoderm invagination, these cells spread out to form a single cell layer and this morphogenetic event is controlled by signaling through the FGF receptor Htl ^20, 28^. Embryos heterozygously mutant for *htl* are viable, but homozygous *htl* mutant embryos exhibit severe mesoderm layer defects and differentiation defects and die during late embryogenesis ^21-23^.

It is impossible to select *htl* homozygous embryos at early stages, because the phenotype of the *htl* mutation cannot be seen under a dissecting microscope. Therefore, to select homozygous *htl* mutant embryos before gastrulation, we made use of a loss of function mutation in the *halo* gene^32^. *halo* mutant embryos are viable and fertile, but exhibit a readily visible mutant phenotype that can be scored in living embryos before gastrulation commences ^30^ (Fig. 2). The *halo* gene is required for proper transport of lipid droplets; in the late blastoderm embryo, lipid droplets are transported from the embryo periphery in a basal direction towards the central yolk. In *halo* mutant embryos this transport is blocked and under transmitted light the periphery of the embryo remains opaque due to the persisting lipid droplets in the embryonic periphery (Fig. 2). This phenotype can be rescued by a transposon insertion containing the genomic *halo* sequence, called *p[halo*^*+*^]^31^. We transposed the *halo* rescue transgene *p[halo*^*+*^] onto balancer chromosomes. Balancer chromosomes are used to maintain recessive mutations in inbred fly lines in a way that all flies are heterozygous for the mutation and for the balancer chromosome. Thus, in a typical inbred cross of heterozygous *htl* parents, one quarter of the embryos are homozygous mutant for the *htl* mutation and do not carry the balancer chromosome. In a *halo* mutant background, the *htl* mutant embryos will have lost the *p[halo*^*+*^] balancer chromosome and therefore represent the only embryos that will show the *halo* phenotype (Fig. 2). For each experiment *halo* mutant embryos from the *halo*^*AJ*^ ; *htl*^*AB42*^ / TM6 *p[halo*^*+*^] stock were staged until they reached mid gastrulation stages (stages 7/8) when they were collected on dry ice to immediately stop development ^39^ (Fig. 2). Obtaining 40 µg of total protein required to collect approximately 100 tightly staged embryos for each biological replicate. Because the *halo* mutation itself could have an effect on the global proteome, we labeled *halo*^*AJ*^ homozygous mutant embryos with Lys-8 as control sample for comparison with *halo*^*AJ*^ ; *htl*^*AB42*^ heterozygous embryos. Additionally, we compared Lys-8 labeled *halo*^*AJ*^ embryos with Lys-0 unlabeled *halo*^*AJ*^ embryos to identify false positive candidates, which might be caused by the stable-isotope labeling itself (see below).

**Figure 2:**
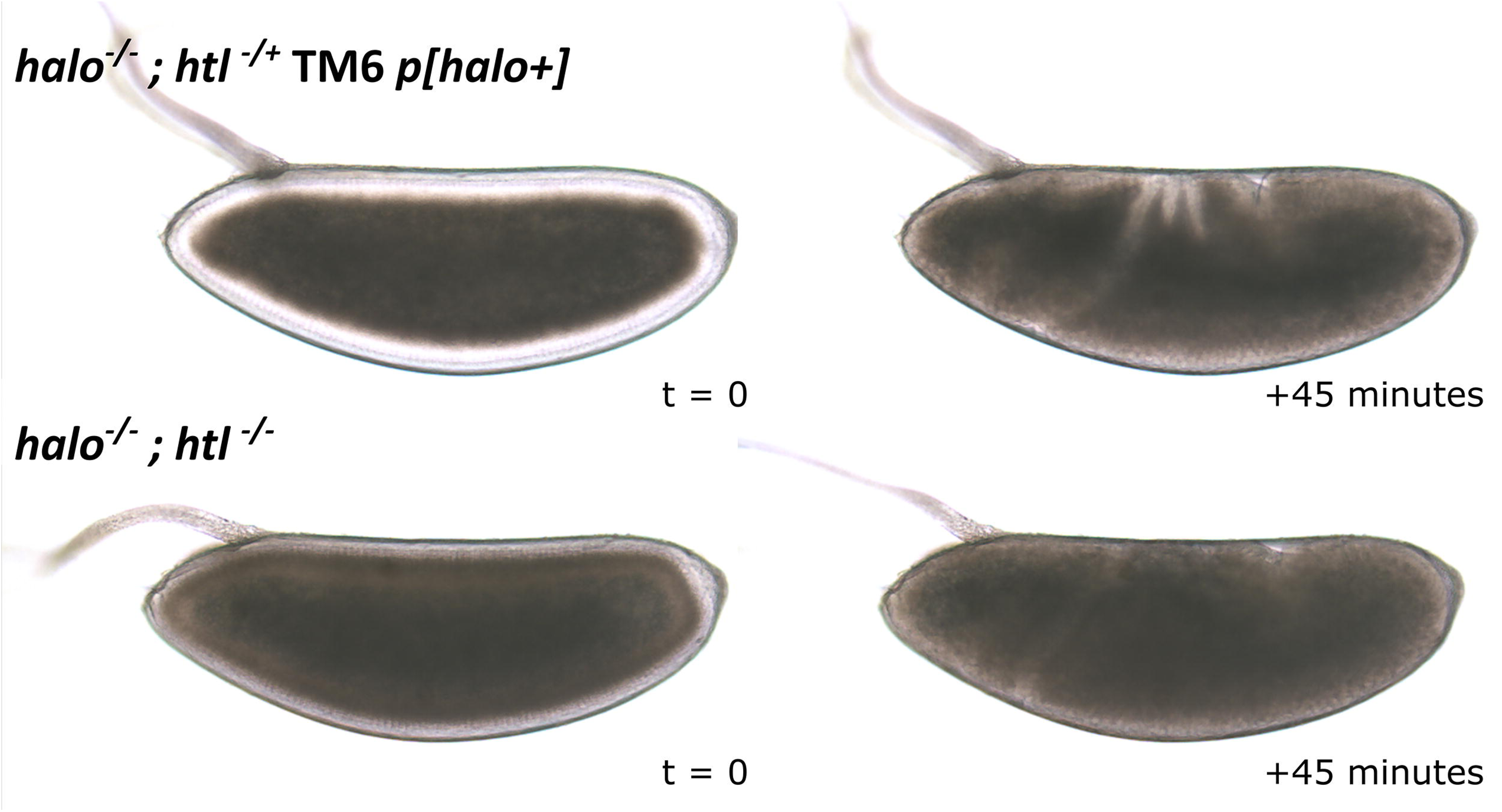
*halo* as a selection marker for the identification of *htl* mutant embryos. The halo *gene* is located on the second chromosome and the *htl* gene is located on the third chromosome. The genomic region containing the *halo* gene was inserted onto the third chromosomal TM6 balancer chromosome. Embryos mutant for *halo*^*AJ*^ develop an opaque ring during cellularization (lower left panel), which is fully rescued by the TM6 *p[halo*^*+*^] balancer chromosome (upper left panel). *heartless*^*AB42*^ embryos were therefore identified by the *halo*^*AJ*^ phenotype, which indicates the absence of the balancer chromosome. The *halo* phenotype is only detectable during stage 5 of embryogenesis and disappears once gastrulation has started (right panel).

### SILAC-based quantitative proteomic analysis of *htl* mutants

Flies that were homozygously mutant for *halo* were labeled with Lys-8 as described in the methods section. We found that a single generation reared on Lys-8 labeled yeast was sufficient to produce isotope-labeled flies. Embryos derived from homozygous *halo*^*AJ*^ SILAC flies showed robust incorporation of Lys-8 at a ratio of over 92% (Fig. 1D). The comparison of Lys-8 labeled embryos of *halo* mutants with Lys-0 unlabeled *halo* mutant embryos did not show any major changes in protein ratios indicating that the stable isotope labeling itself did not produce false positives (Fig. S2). For quantitative proteomic analysis, we collected late gastrula embryos from a Lys-8 labeled homozygous *halo*^*AJ*^ ; *htl*^*+*^ stock and Lys-0 labeled homozygous *halo*^*AJ*^ ; *htl*^*AB42*^ embryos (Figs. 1, 2). 40 µg of total protein in each experiment and 3 replicates were analyzed independently via LC-MS/MS. Overall, the experiments detected a total number of 81.719 peptides including the comparison of the Lys-8 vs. Lys-0 labeled *halo*^*AJ*^ experiment. Using MaxQuant analysis these peptides were assigned to 2.131 proteins. In between the three experimental repliates 994 proteins were found in all the experiments and therefore were selected for further analysis. The average sequence coverage of these proteins was at 24.57% (STDEV = 15.92%). First, we tested the overall changes in protein abundance between embryos derived from *halo* control and *htl* mutant flies. The correlations of the three biological replicates with each other was tested in pairs by plotting the Log 2 ratios of heavy to light labeled proteins using Perseus ^35^. The population histograms showed a positive correlation of the ratios of heavy to light labeled proteins between all three bio replicates (Fig. 3). Scatter plot analyses of the individual replicates to each other confirmed a positive relationship with Pearson correlation co-efficient values larger than 0.6, indicating that the biological replicates indeed produced a large degree of overlapping data points (Fig. 3).

**Figure 3:**
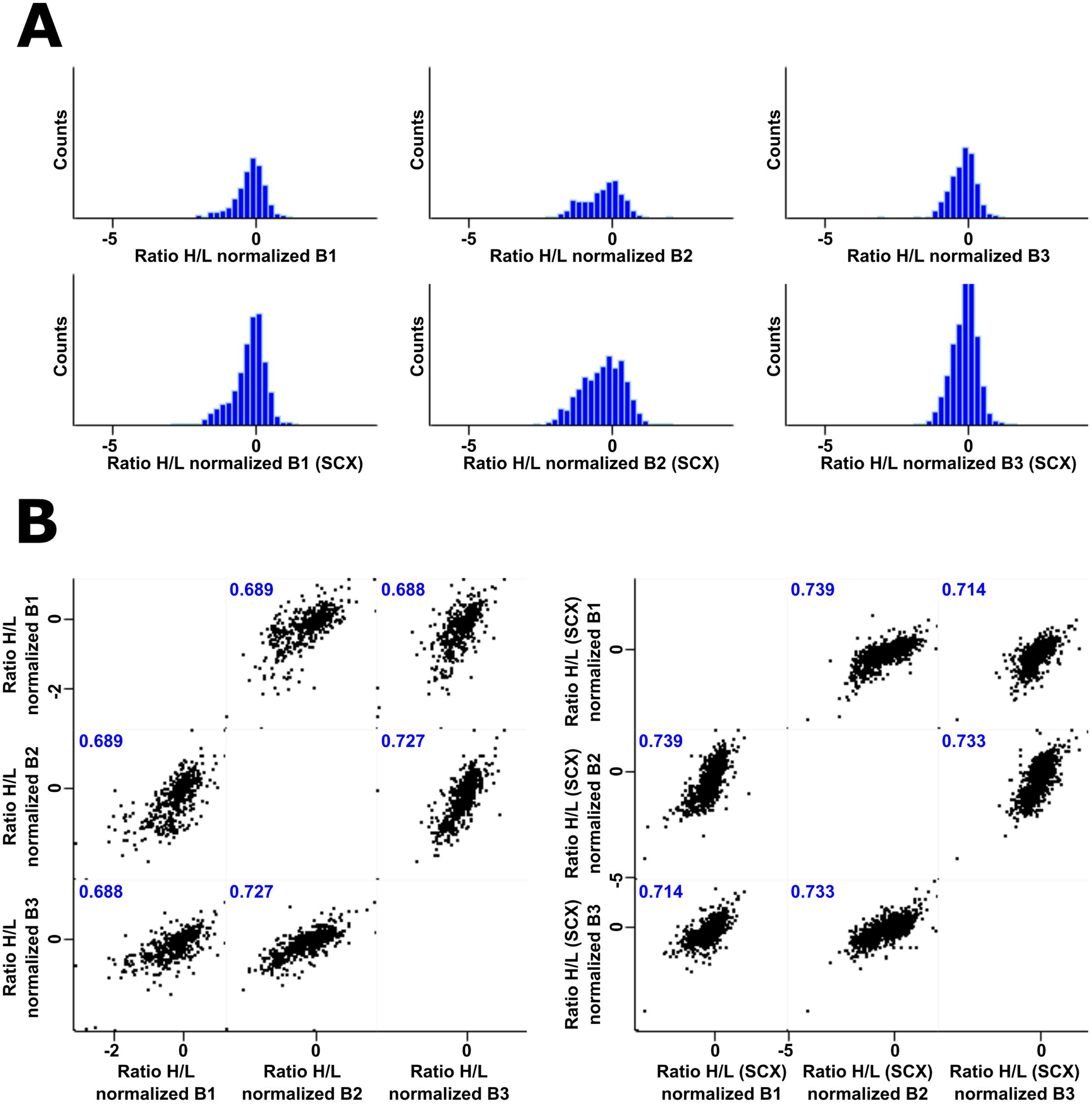
Population statistics of *htl*^*AB42*^ embryos. **(A)** Lys-8 (H) over Lys-0 (L) protein ratios were found to be normally distributed in all 3 biological replicates. Upper panel shows H/L ratios of unenriched sample sets and the lower panel H/L ratios of the respective SCX enrichments. H/L protein ratios represent normalized values. **(B)** Scatter plots indicate correlation of the individual biological replicates to each other with Pearson correlation co-efficient larger than 0.6. Left scatter plot represents comparison of unenriched data sets, whereas the right one shows correlation after SCX enrichment.

### Changes of protein abundance in *htl* mutant embryos

The positive correlation of the replicates provided a basis to determine changes in protein abundance that were consistent in all three experimental repeats. The median and standard deviations of the population distributions were calculated to determine the average cut-off values for up- and down-regulated proteins within fold-change range of +/- 1.1. We found that 36 proteins were consistently down-regulated in *htl* mutant embryos, whereas 25 proteins were consistently up-regulated in all three biological replicates (Table 1, Table 2). We also detected six significant changes (two up-regulated and four down-regulated) in protein regulation when comparing Lys-8 labeled *halo* mutant embryos with Lys-0 unlabeled *halo* mutant embryos. These changes were scored to be false positive identifications and were not considered in further analyses (Fig. S2B). The positive correlation between the three bio replicates allowed for the examination of statistically relevant changes in the protein abundances between control and *htl*^*AB42*^ mutant embryos. To consider both the degree of fold changes of proteins and the statistical significance (-logP value) of the change, we visualised the data using a Volcano plot (Fig. 4).

**Table 1:**
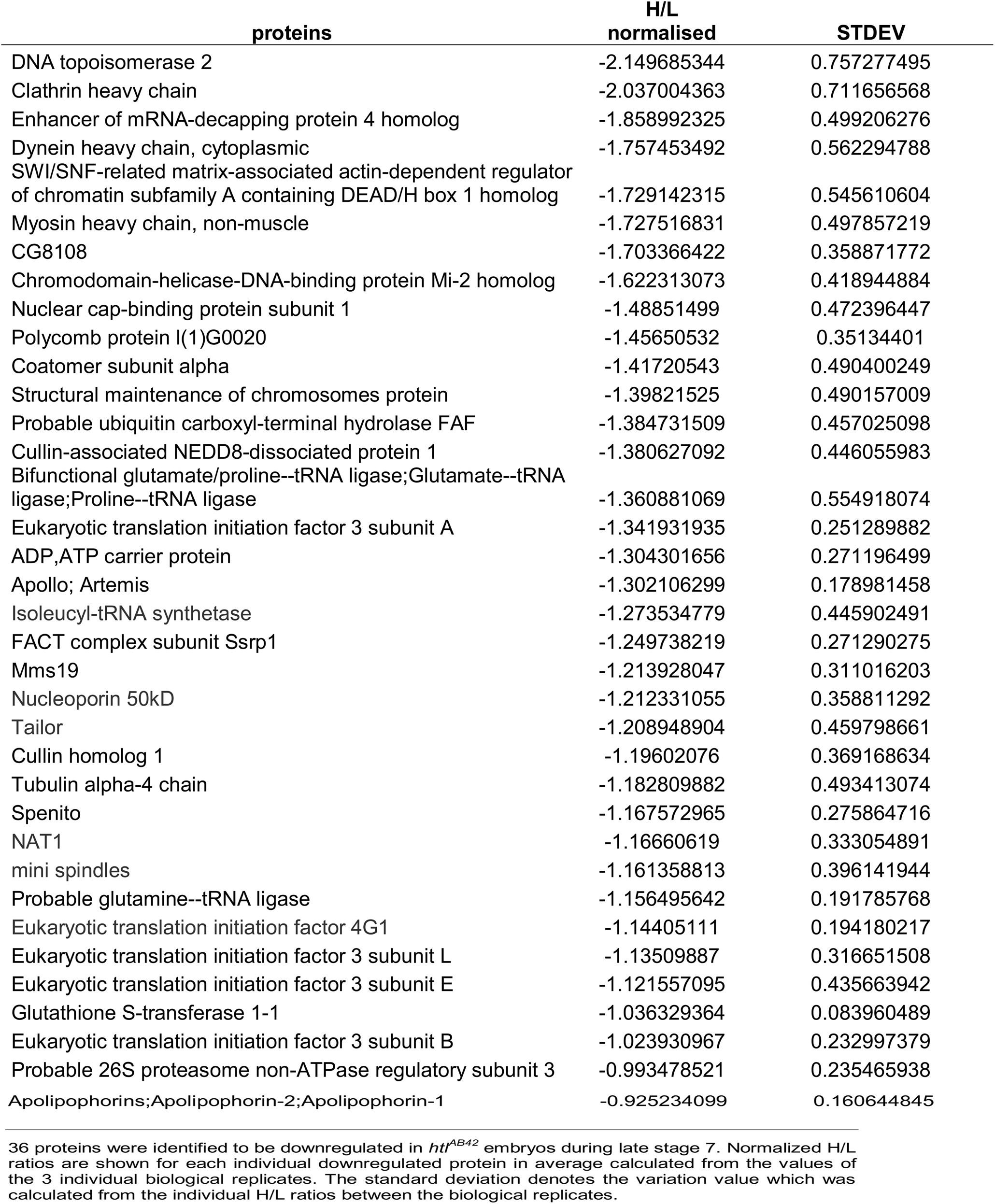
Proteins downregulated in homozygous *htl* mutant embryos.

**Table 2:** Proteins upregulated in *htl* homozygous embryos.

| Up-regulated proteins | H/L normalised | STDEV |
| --- | --- | --- |
| Asparagine synthetase | 1.607497445 | 0.048613934 |
| Dipeptidase B | 1.586862616 | 0.324155904 |
| Peptidyl-prolyl cis-trans isomerase | 1.289350727 | 0.087768972 |
| Glutathione S-transferase S1 | 1.147126148 | 0.29662771 |
| CG6084 | 1.116092892 | 0.105111356 |
| Dihydropteridine reductase | 1.002222371 | 0.147232873 |
| Succinyl-CoA:3-ketoacid-coenzyme A transferase | 0.93689031 | 0.145506671 |
| Probable histone-binding protein Caf1 | 0.910630778 | 0.293233751 |
| CTP synthase | 0.782985113 | 0.249967156 |
| CG2915 | 0.780524633 | 0.329577881 |
| Maltase A5 | 0.778792229 | 0.207563376 |
| Nucleoplasmin-like protein | 0.775825308 | 0.222484798 |
| Aldehyde dehydrogenase | 0.734156969 | 0.186440997 |
| Ecdysone-induced protein 55E | 0.712951314 | 0.254806708 |
| Glutathione S transferase E13 | 0.700371724 | 0.053918198 |
| Microsomal glutathione S-transferase-like | 0.679973556 | 0.109014319 |
| CG4069 | 0.67469731 | 0.158609716 |
| CG17337 | 0.67256631 | 0.259314925 |
| CG9149 | 0.668636923 | 0.059261456 |
| Aldehyde oxidase 1 | 0.666252533 | 0.229121238 |
| vibrator | 0.637438132 | 0.158088087 |
| alphabet | 0.614572157 | 0.11629388 |
| Peptidyl-prolyl cis-trans isomerase;12 kDa FK506-binding protein | 0.581425911 | 0.070975358 |
| CG6028 | 0.574864157 | 0.095288709 |
| CG6745 | 0.563803843 | 0.003837915 |

25 proteins were identified to be up-regulated in *htl*<sup>AB42</sup> embryos during late stage 7. Normalized H/L ratios are shown for each individual up-regulated protein in average calculated from the values of the 3 individual biological replicates. The standard deviation denotes the variation value calculated from the individual H/L ratios between the biological replicates.

**Table 3:**
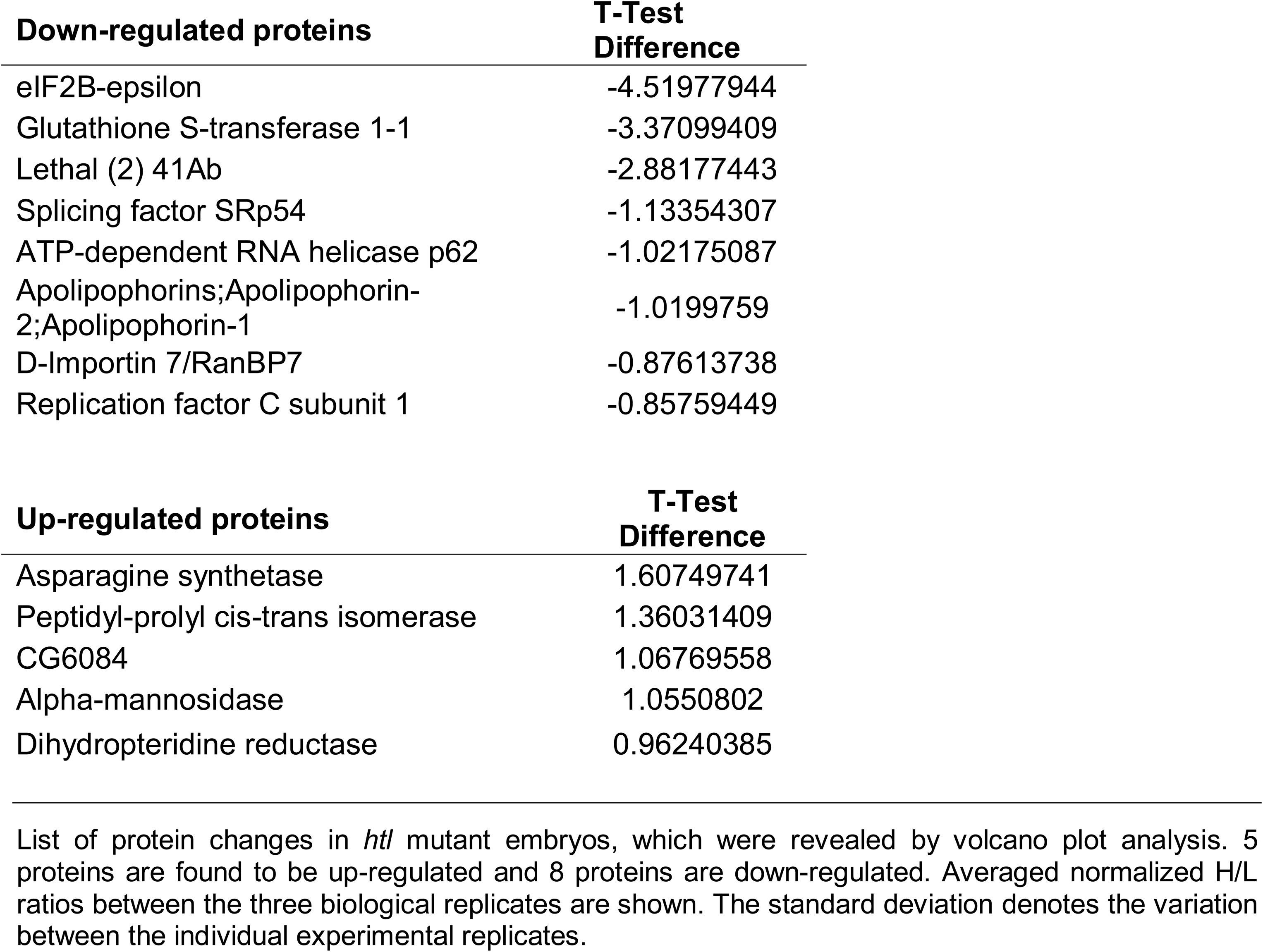
candidates of down- and upregulated proteins according to statistical evidence.

**Figure 4:**
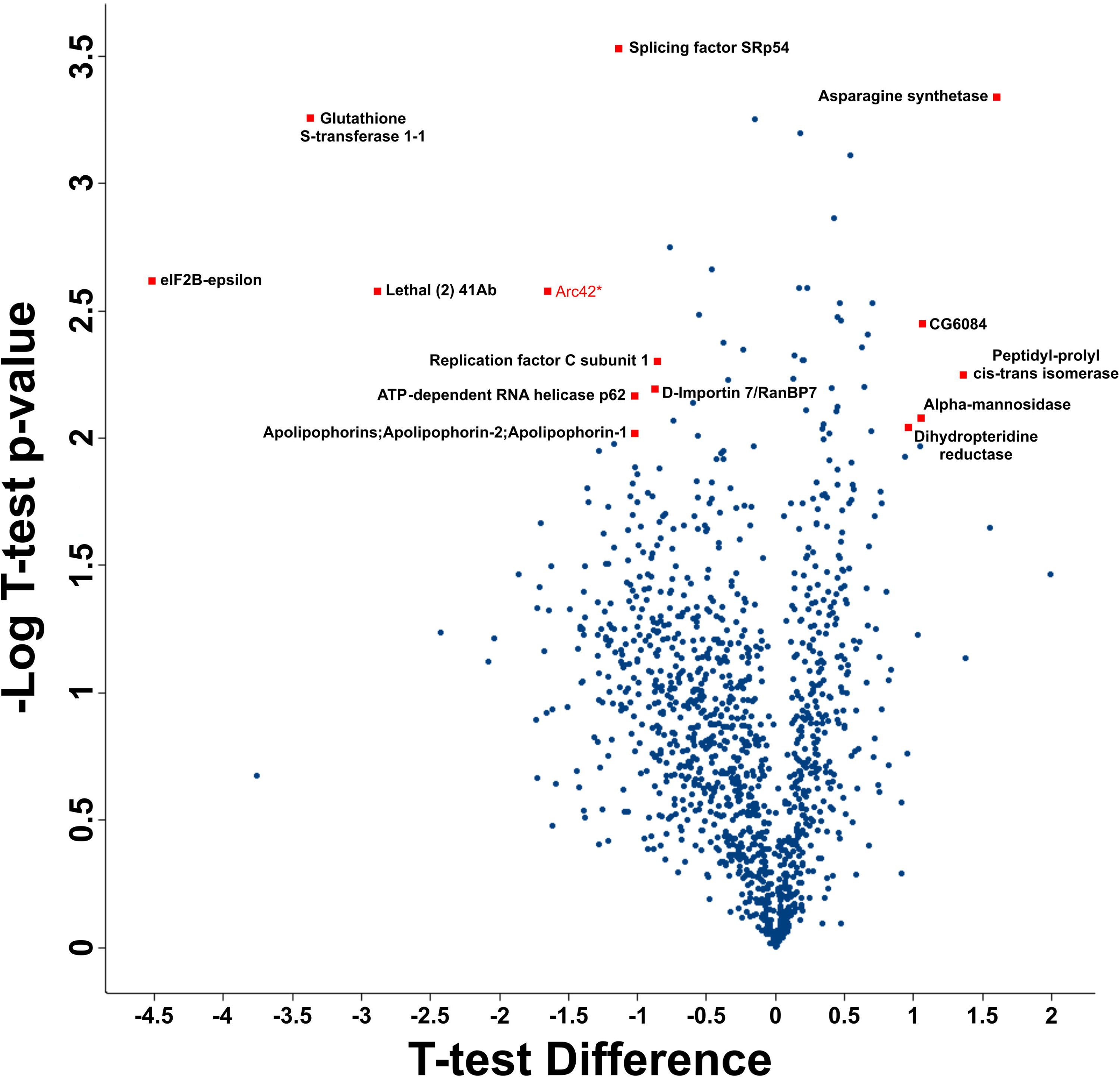
Changes in the proteome of FGF Htl deficient embryos. Vulcano plot depicting the quantification of proteome changes of *htl* mutant embryos during gastrulation. Relative fold changes are depicted as log2 averaged SILAC ratios of all three biological replicates and were plotted against the -log10 t-test p-value. Up and down regulated proteins are indicated with cut-offs +/-1. One particular candidate (Q9VDT1, Arc42) was also identified as down-regulated when comparing Lys-8 *halo* vs. Lys-0 *halo* mutants and thus considered as false positive.

To determine whether the down- or up-regulated proteins exhibited linked functional features, we performed a String network analyses^36^. These analyses revealed that 31 of the down-regulated proteins and 18 of the up-regulated proteins were linked and belonged to networks for which String found independent evidence of interaction. In *htl* mutant embryos the largest class of down-regulated proteins was associated with chromatin (Fig. 5). Other down-regulated proteins were found to occur in networks that included intracellular transport, mRNA binding/processing and translation. Interestingly, we found central components of the endomembrane transport machinery, including Clathrin heavy chain, Vps35, and the coatomer component COP1 alpha. In addition, we also found cytoskeletal components like myosin heavy chain, Tubulin, the microtubule regulator Mini spindles and Dynein heavy chain 64C to be down-regulated in *htl*^*AB42*^ mutant embryos. To a smaller extent we detected down-regulation of some metabolic and cytoskeletal components in *htl*^*AB42*^ embryos. In contrast to the downregulated proteins, the largest network detected to be up-regulated in *htl*^*AB42*^ embryos is affecting metabolic pathways. Some proteins affecting chromatin and cytoskeletal networks were additionally found to be up-regulated. A small number of proteins could not be assigned to particular networks (Figs. 5,6).

**Figure 5:**
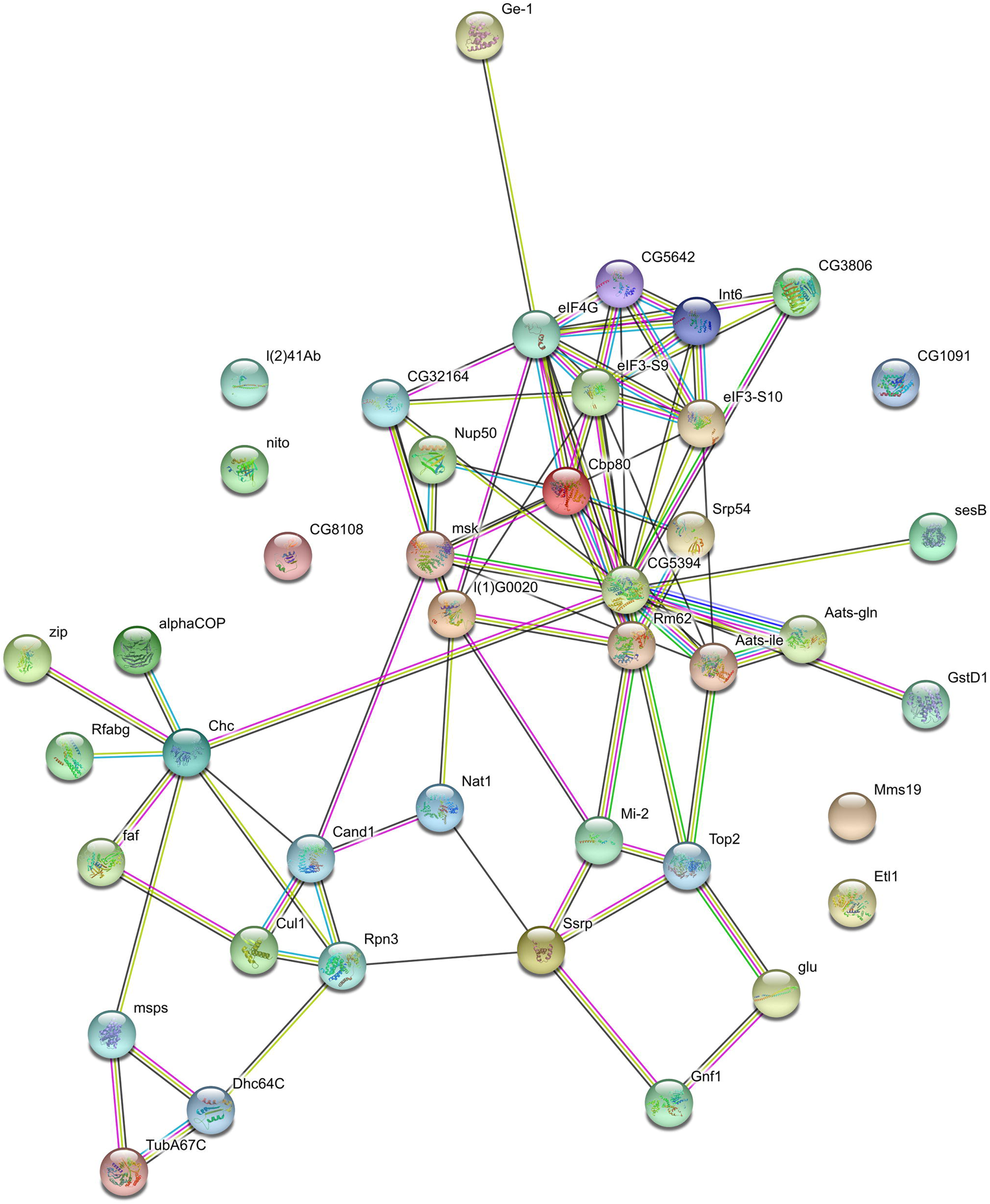
String analysis of down-regulated proteins. Summary view of the network of down-regulated proteins in embryos lacking the Htl FGF signaling pathway during gastrulation. The network analysis indicates the down-regulation of proteins that are associated with chromatin, nuclear transport, the binding or processing of mRNA, and translation. Central components of the cytoskeleton, such as myosin heavy chain, Tubulin, the microtubule regulating protein Mini spindles and Dynein heavy chain 64C, are also down-regulated when Htl FGF signaling is depleted. A few down-regulated components belong to metabolic and cytoskeletal pathways but could not be assigned to any particular network in *htl* mutant embryos.

**Figure 6:**
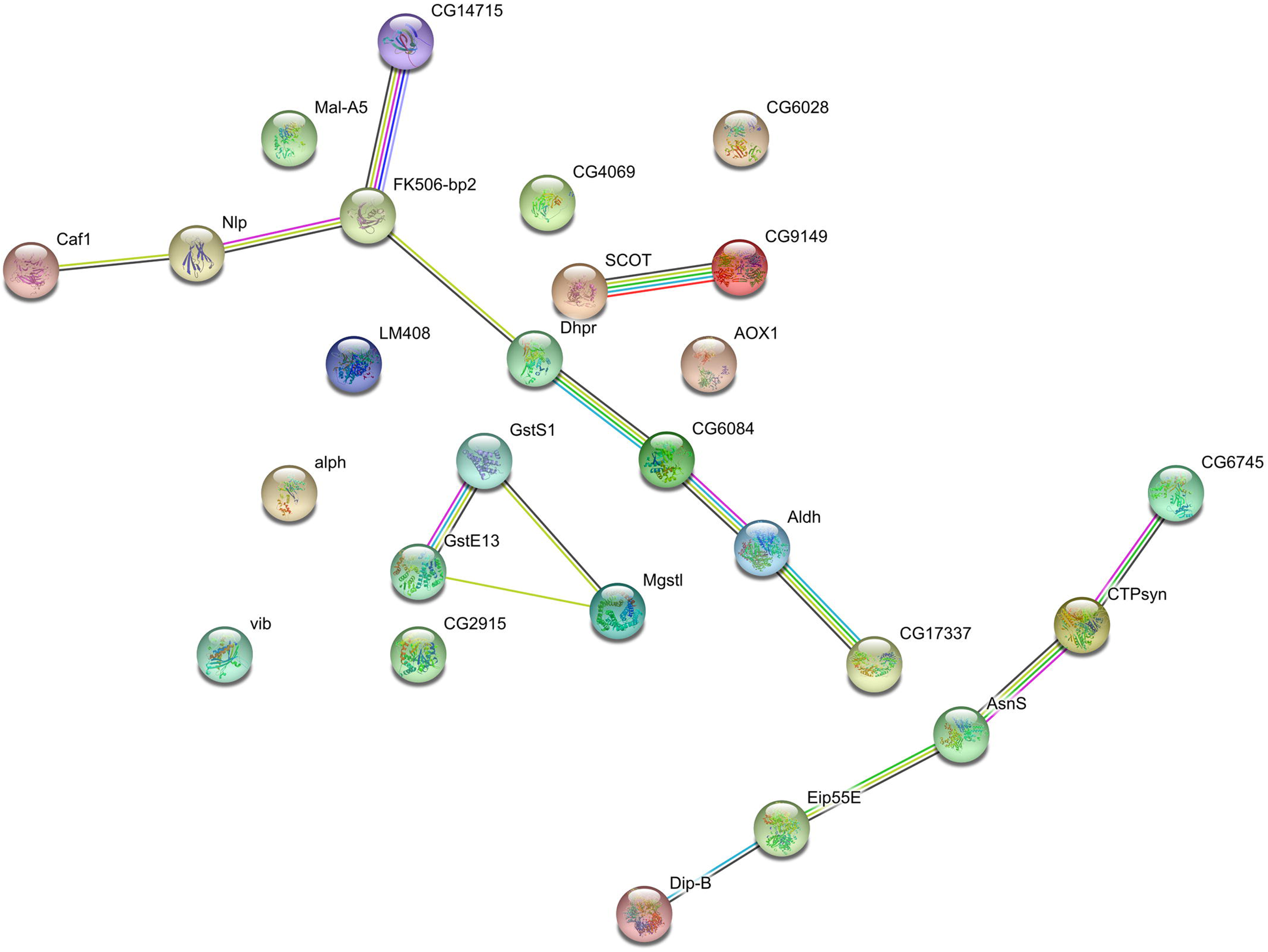
String analysis of up-regulated proteins. Summary view depicting the network of up-regulated proteins in embryos lacking Htl FGF signaling. Most of the identified up-regulated proteins were assigned to metabolic pathways, whereas some proteins are belonging to chromatin and cytoskeletal networks. A few candidates were not assigned to any particular network.

## Discussion

Although *Drosophila* is one of the most studied model organisms, quantitative proteomics have surprisingly not been applied extensively for the analyses of embryonic mutants ^9, 11, 12^. One possible reason for that might be caused by problems arising when growing healthy populations of flies with incorporated stable non-radioactive isotopes. Furthermore, *Drosophila* is routinely marked with stable isotope labeled L-Lysine and peptides are therefore generated by lysyl-endopeptidase treatment. This endopeptidase is required to ensure all peptides generated contain at least one labeled amino acid, resulting in Lys8-labeled peptides being larger compared to tryptic peptides, which arise from cleavage at Lysine and Arginine sites, thereby resulting in reduced sequence coverage. Flies survive on an Arginine deficient diet, thus reducing its usefulness as a source for stable isotope labeling of proteins ^40^. Furthermore, it has been reported that both Lysine and Arginine are metabolized into several other amino acids in the fly and this can affect the quantification ^16^. Genome editing of enzymes involved in Arginine synthesis may improve the use of stable isotope labeled Arginine as a second label.

In this work, we were able to conduct a global proteomic analysis of embryos depleted for Htl FGF receptor signaling by combining genetics with a modified, more feasible and cost-efficient way of labeling *Drosophila* with stable non-radioactive heavy L-Lysine. The SILAC fly was established previously, however unfortunately, we were not able to obtain large enough quantities of flies following these protocols confirming reports indicating growth retardation and low survival rates of larvae by replacing normal fly food with stable isotope labeled yeast ^9, 11^. Here we introduce a protocol that overcomes the decreased fitness of larvae and flies in these procedures and produces robustly labeled *Drosophila* embryos in a single generation. We found that animals raised on apple juice agar supplied with Lys-8 labeled yeast strain *BY4742*, were eclosing with expected ratios and appeared healthy. Our results suggest that low eclosing rates that we were observed using other protocols might due to the minimal food and lack of minerals and/or vitamins. We conclude that the present protocol produces robustly labeled embryos while requiring only low amounts of labeled yeast, which together makes the SILAC fly an economically attractive approach.

Despite the feasibility of the protocol for labeling *Drosophila* with stable non-radioactive isotopes, we came across some limitations for quantitatively studying the global proteome. A single embryo only contains around 1 µg of protein, which requires to collect a large number of staged embryos for quantitative global analyses. In our experimental setup we did not detect many phosphorylated peptides, which can be explained by the low amount of protein that we have used for each replicate. To extend the global proteomic analysis of FGF receptor-depleted embryos towards the phospho-proteome will require to scale up each biological replicate by a factor of 20 in order to obtain enough material for enrichment of phospho-peptides using TiO_2_ or Ti-IMAC ^41, 42^. Alternatively, large-scale quantification of phosphorylated peptides can be also combined by the spike-in SILAC method ^43, 44^.

Since we wanted to monitor the proteomic change in embryos depleted for FGF signaling during gastrulation stages, it was necessary to establish a good and reliable marker for the selection of embryos homozygous for the mutation in the FGF receptor Htl. Here we established a method using the *halo* mutation as a genetic marker, which is readily visible in transmitted light in live embryos ^29^. Linking a transgenic *halo* rescue construct with a balancer chromosome allowed us to reliably select *htl* homozygously mutant embryos before gastrulation by the presence of the *halo* phenotype. Other methods like, using GFP linked to balancer chromosomes, are not able to provide a reliable signal to noise ratio for efficient selection under a dissecting microscope. *halo* has been previously used for early selection of homozygous mutations, but its use was restricted for genes located on the second chromosome ^32^. The establishment of balancers containing the transgenic *p[halo*^*+*^] in a *halo* mutant background open the opportunity for the application of this technique to other experiments in order to genotype and select embryos homozygous for zygotic mutations before the actual phenotype occurs.

The *halo* linkage method was employed to collect tightly staged, homozygously *htl* mutant gastrula embryos for quantitative proteome analyses. By LC/MS-MS we compared unlabeled *htl* mutant embryos with stable isotope (L-Lysine ^13^C_6_ ^15^N_2_) labelled *halo* embryos as control. Among 2131 identified proteins in all three biological replicates we identified 36 proteins to be down-regulated, whereas 25 were found to be up-regulated. Our data indicate that the lack of Htl FGF receptor signaling in the early embryo affects the abundance of proteins involved in the regulation of chromatin, nuclear transport, mRNA function, and endomembrane transport as well as the cytoskeleton. The majority of up-regulated proteins are related to various metabolic pathways including amino acid biosynthesis, carbohydrate metabolism. Among proteins which levels change, we only detected one protein that was known to be directly involved in the FGF signaling pathway in *Drosophila*. We found the Drosophila homolog of Importin 7, encoded by the gene *moleskin* (*msk*), to be downregulated in the mutants. Interestingly, Msk has been shown to be required for FGF signaling through the Breathless (Btl) receptor in tracheal development ^45^. The functions of Msk include mediating the nuclear transport of activated ERK and may also be related to its role in promoting activation of Rac signaling, which has also been demonstrated to be critical for mesoderm layer formation^45-47^.

The most interesting class of proteins were those, which levels were reduced in embryos lacking Htl signaling, because they represent candidates for a function or a response that is dependent on FGF-dependent signaling events that cause morphogenetic changes. Consistent with an important function of the Htl FGF receptor in triggering cell migration and protrusion formation, we found that the levels of tubulin 67A and the microtubule plus end binding protein Mini spindles, cytoplasmic dynein heavy chain 64C and non-muscle myosin heavy chain were reduced in the mutants. The problably most provocative group of candidates were proteins involved in intracellular transport processes including endomembrane transport (Clathrin heavy chain and COP1 alpha). The membrane transport proteins are components of two distinct transport processes; Clathrin mediates transport in the endosomal system while COP1alpha coated vesicles are involved in secretory pathway and retrograde transport within the Golgi complex^48^. Binding of FGF ligand to its receptor stimulates Clathrin-dependent receptor endocytosis, which has been considered both as one mechanism of signal attenuation, but also as a mechanism of signal propagation^49^. In the case of COP1 alpha it is interesting to note that assembly of the COP1 coatomer requires the small GTPase ARF1, which plays a crucial role in protrusion formation during cell migration^50, 51^. Further studies will require whether FGF signaling impacts on nuclear transport (Moleskin and Artemis/Apollo1 [RanGTP binding proteins, and Nup50]) and protein turnover (Cullin1, Cand1 and Rpn3). The discovery of changes in amounts of interesting proteins that fall into functionally related classes presents a starting point for further functional analyses. The most important issue will be to determine the tissue-specificity of potential protein functions in FGF-receptor induced responses. *Drosophila* provides a rich resource to tackle this problem, for example by tissue-specific RNAi-mediated gene knock-down experiments^52^ or tissue-specific protein knock down^53^.

## Acknowledgements

We thank Michael Welte (Univ. Rochester, U.S.A) for the gift of the *halo* fly stocks and discussions. We thank Angus Lamond (University of Dundee, U.K.) and Matthias Trost (University of Newcastle, U.K.) for support and discussions throughout this study. We thank Ryan Webster (University of Dundee, U.K. for expert technical assistance and Kelly Hodge for assistance with mass spectrometry, Elham Gheisari for help with preparing the figures and Katja Kapp for critical comments to the manuscript. Stocks obtained from the Bloomington Drosophila Stock Center (NIH P40OD018537) were used in this study. This study was funded by MRC project grant KO18531/1 to HAJM and a University of Kassel stipend ‘future programme leader award’ to HB.

## Figures legends

**Figure S1: Incorporation of Lys-8 in yeast *SUB62 vs. BY4742* strains**

**(A,B)** Label check on *SUB62* grown in 1l Lys-8 containing culture. Global peptide ratios **(A)** and representative spectra **(B)** from *SUB62* grown to saturation in 1l Lys-8 culture. An average global SILAC ratio **(A)** of 0.980 indicates incomplete SILAC labeling of *SUB62*, **(B)** representative spectra of FALGQGVGVILCIGETLEEK peptide from Triosephosphate isomerise, with a calculated SILAC ratio of 1.110. **(C**,**D)** Label check on *BY4742* grown in 1l Lys-8 culture. Global peptide ratios **(C)** and representative spectra **(D)** from *BY4742* grown to saturation in 1l culture supplemented with Lys-8. An average global SILAC ratio **(C)** of 14.912 indicated complete SILAC labeling of *BY4742*. **(D)** Representative spectra of SRSGVAVADESLTAFNDLK peptide from COF1p, with a calculated SILAC ratio of 15.585. **(E)** *SUB62* contains a revertible *lys2* mutation. Growth of *SUB62* **(E’)** and *BY4742* **(E’’)** strains on complete media plates (left panel) and complete media plates lacking Lysine (right panel). **(E’’)** *BY4742* did not grow on plates lacking Lysine when plated from frozen stock, or from 1l culture. **(E’)** Growth of *SUB62* on plates lacking Lysine was observed after growth in 1l culture, but not when plated from frozen stock.

**Figure S2: Comparison of Lys-8 labeled *halo*^*AJ*-/-^ with Lys-0 unlabeled *halo*^*AJ*-/-^ embryos.** Four proteins were found to be down-regulated when comparing Lys-8 vs. Lys-0 labeled *halo* mutants, whereas two proteins were found to be up-regulated. Respective proteins were removed from the candiates which were identified to be up- and down-regulated comparing the three biological replicates.

