## Supplementary figures and images for "SILAC-based quantitative proteomic analysis of *Drosophila* gastrula stage embryos mutant for fibroblast growth factor signaling"

### Fig. S1

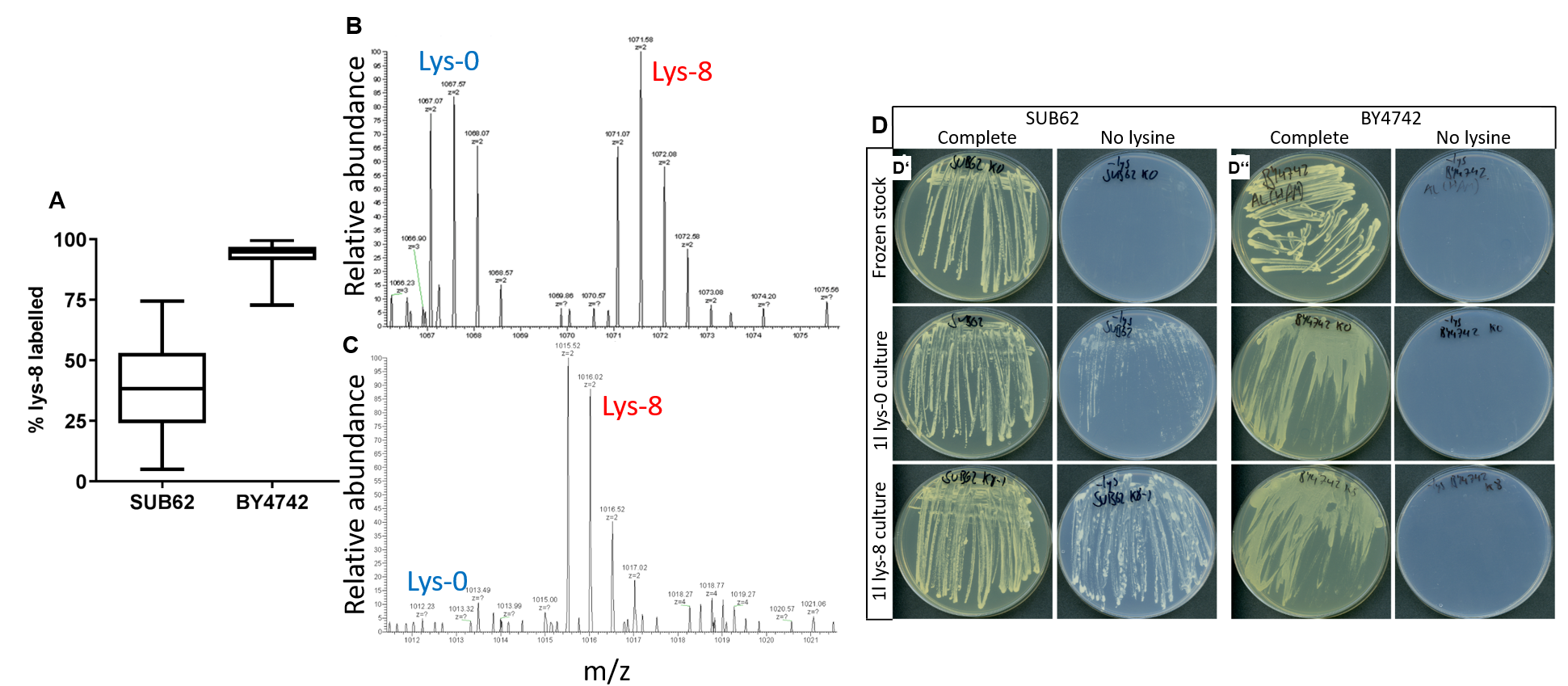

### FIg. S2

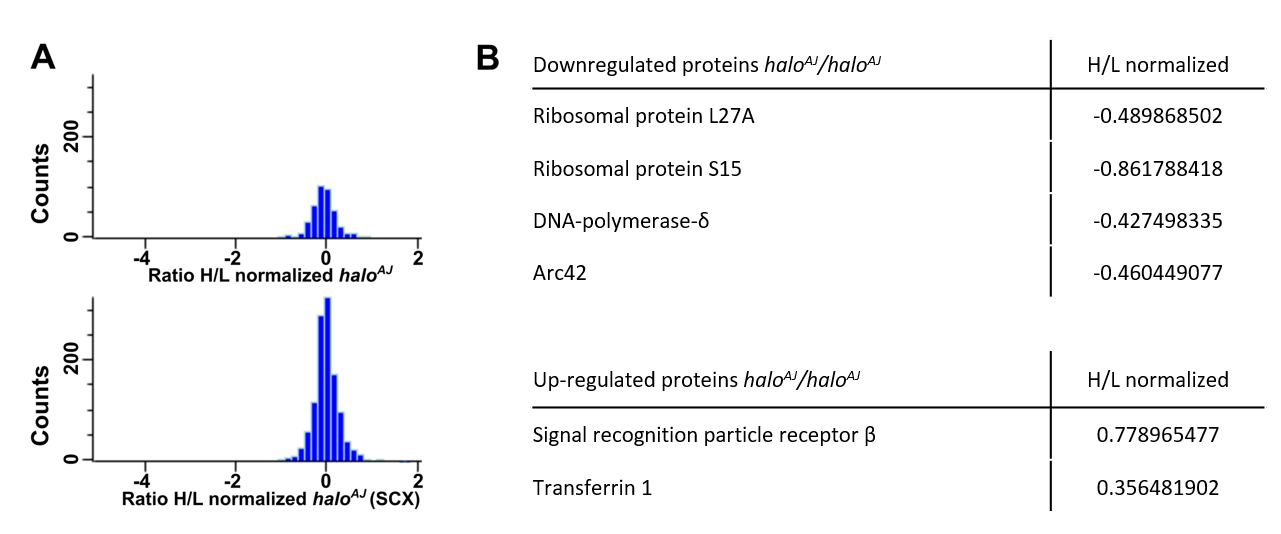
